# Mucosal CD8^+^ T cell responses induced by an MCMV based vaccine vector confer protection against influenza challenge

**DOI:** 10.1101/494864

**Authors:** Xiaoyan Zheng, Jennifer D. Oduro, Julia D. Boehme, Lisa Borkner, Thomas Ebensen, Ulrike Heise, Marcus Gereke, Marina C. Pils, Astrid Krmpotic, Carlos A. Guzmán, Dunja Bruder, Luka Čičin-Šain

## Abstract

Cytomegalovirus (CMV) is a ubiquitous β-herpesvirus that establishes life-long latent infection in a high percentage of the population worldwide. CMV induces the strongest and most durable CD8^+^ T cell response known in human clinical medicine. Due to its unique properties, the virus represents a promising candidate vaccine vector for the induction of persistent cellular immunity. To take advantage of this, we constructed a recombinant murine CMV (MCMV) expressing an MHC-I restricted epitope from influenza A virus (IAV) H1N1 within the immediate early 2 (*ie2*) gene. Only mice that were immunized intranasally (i.n.) were capable of controlling IAV infection, despite the greater potency of the intraperitoneally (i.p.) vaccination in inducing a systemic IAV-specific CD8^+^ T cell response. The protective capacity of the i.n. immunization was associated with its ability to induce IAV-specific tissue-resident memory CD8^+^ T (CD8T_RM_) cells in the lungs. Our data demonstrate that the protective effect exerted by the i.n. immunization was critically mediated by antigen-specific CD8^+^ T cells. CD8T_RM_ cells promoted the induction of IFNγ and chemokines that facilitate the recruitment of antigen-specific CD8^+^ T cells to the lungs. Overall, our results showed that locally applied MCMV vectors could induce mucosal immunity at sites of entry, providing superior immune protection against respiratory infections.

**Author summary:** Vaccines against influenza typically induce immune responses based on antibodies, small molecules that recognize the virus particles outside of cells and neutralize them before they infects a cell. However, influenza rapidly evolves, escaping immune recognition, and the fastest evolution is seen in the part of the virus that is recognized by antibodies. Therefore, every year we are confronted with new flu strains that are not recognized by our antibodies against the strains from previous years. The other branch of the immune system is made of killer T cells, which recognize infected cells and target them for killing. Influenza does not rapidly evolve to escape T cell killing; thus, vaccines inducing T-cell responses to influenza might provide long-term protection. We introduced antigen from influenza into the murine cytomegalovirus (MCMV) and used it as a vaccine vector inducing Killer T-cell responses of unparalleled strength. Our vector controlled flu replication and provided relief to infected mice, but only if we administered it through the nose, to activate killer T cells that will persist in the lungs close to the airways. Therefore, our data show that the subset of lung-resident killer T cells is sufficient to protect against influenza.

## Introduction

Respiratory infections caused by influenza viruses usually are associated with mild-to-moderate disease symptoms but are linked with high morbidity and mortality in susceptible populations like the elderly, young children, patients with co-morbidities and immunocompromised patients. Influenza virus causes seasonal epidemics, with typically 3 to 5 million cases of severe illness worldwide, according to WHO reports [1], and influenza type A viruses (IAV) cause the more severe disease form. Vaccines against influenza are based on the induction of adaptive immunity that targets the projected yearly epidemics. While most vaccines are based on inactivated IAV formulations inducing anti-IAV IgG responses, live attenuated influenza vaccines (LAIV) are also used as formulations for i.n. administration. This is based on the assumption that the induction of local immunity may provide superior immune protection [2, 3]. However, it remains unclear whether this protection depends on local IgA responses, on cytotoxic T cell responses, or on their combined antiviral activity. Of note, functional T cell responses were shown to substantially contribute to antiviral IAV immunity [4–6]. In particular, cytotoxic influenza-specific CD8^+^ T lymphocytes (CTLs) promote viral clearance indirectly by secretion of pro-inflammatory cytokines such as IFNγ [7] and directly by perforin/Fas-mediated killing of infected epithelial cells in the bronchoalveolar space [8]. However, it remained unclear if T cell responses alone may control IAV, or if Ig responses were the crucial contributor to LAIV-mediated immune protection. We considered that this question could be addressed by developing a vaccine formulation that optimizes T cell responses against IAV, while excluding the humoral ones.

CMV infection induces sustained functional T cell responses that are stronger in the long-term than the immune response to any other infectious pathogen [9]. Experiments in the mouse model have shown that defined CMV epitope-specific CD8^+^ T cells accumulate in tissues and blood and are maintained at stable high levels during mouse CMV (MCMV) latency [10]. This phenomenon was termed ‘‘Memory Inflation’’ [11]. While some MCMV derived peptides, as the ones derived from the IE3 (**_416_RALEYKNL_423_**) and M139 proteins (**_419_TVYGFCLL_426_**) induce inflationary responses, other peptides, such as the M45-derived (**_985_HGIRNASFI_993_**), induce conventional CD8^+^ T cell response [12]. The exceptionally long-lasting cellular immunity to CMV antigens has raised the interest in CMV as a potential new vaccine vector [13]. Antigen-experienced CD8^+^ T cells are subdivided into CD62L^-^ effector memory (CD8T_EM_) and CD62L^+^ central memory CD8^+^ T cells (CD8T_CM_). The antigen-specific CD8^+^ T cells during latent infection bear predominantly a CD8T_EM_ phenotype and localize in secondary lymphoid or non-lymphoid organs [14]. They may provide immune protection against diverse viral targets [13, 15–18], but also against bacterial [19] or tumor antigens [20, 21].

Both CD8T_EM_ and CD8T_CM_ subsets recirculate between the blood, the lymphoid organs, and the peripheral tissues. A special subset of memory CD8^+^ T cells (CD8T_RM_) resides in non-lymphoid tissues such as lungs, the female reproductive tract (FRT), the skin, the brain or the small intestine [22–25]. These cells lose the capacity of recirculating, maintain themselves at the site of infection, and their phenotype and transcriptional profile differ from classical memory T cells [26]. The well-characterized CD8T_RM_ cells express C-type lectin CD69 [22] and the integrin αEβ7, also known as CD103 [26]. They provide rapid and superior protection against pathogens at the site of infection [22, 26, 27]. A recent publication argued that a vaccine formulation adjuvanted by IL-1β enhances the immune control of IAV by improving mucosal T cell responses [28], but IL-1β improved both humoral and cellular responses in their study. Hence, the contribution of CD8T_RM_ to IAV immune control remains unclear.

CD8T_RM_ are found in the salivary glands of MCMV-infected animals [29], but not in their lungs [30]. We showed that i.n. infection with MCMV induces inflationary CD8^+^ T cell responses, but also that memory inflation is stronger upon i.p. infection [31]. The i.n. administration of an MCMV vaccine vector induced CD8T_RM_ responses in the lungs [25] and only i.n. immunization restricted the replication of respiratory syncytial virus (RSV) upon challenge [25], indicating that CD8T_RM_ elicited by i.n. administration of MCMV vectors might provide immune protection against respiratory virus infections, yet this evidence remains correlative. Upon antigen encounter, CD8T_RM_ cells quickly reactivate at the mucosal site and secrete cytokines and chemokines or support the release of inflammatory mediators by other immune cells [24, 32, 33]. Lung airway CD8T_RM_ cells provide protection against respiratory virus infection through IFNγ and help to recruit circulating memory CD8^+^ T cells to the site of infection in an IFNγ-dependent way [32]. Therefore, to understand if CD8T_RM_ cell may provide immune control of respiratory infections may help to refine strategies for tissue-targeted vaccine design.

In this study, we constructed an MCMV vector expressing the MHC-I restricted peptide **_533_IYSTVASSL_541_ (IVL_533-541_)** [34] from IAV H1N1 hemagglutinin (HA)-MCMV^IVL^ under the transcriptional control of the *ie2* promotor. We investigated the potential of this recombinant virus to induce HA-specific CD8^+^ T cells that confer protection against a lethal IAV challenge. We showed that i.n., but not i.p. immunization with MCMV^IVL^ resulted in a robust protection against an IAV challenge. Protection following i.n. MCMV^IVL^ immunization was associated with high levels of antigen-specific CD8T_RM_ cells in the lungs, and targeted depletion of lung-CD8T_RM_ cells revealed that the control of the IAV in the lungs depended on these cells.

## Results

### Generation of recombinant MCMV and its replication *in vitro* and *in vivo*

We showed recently that MCMV vector expressing a single, optimally positioned MHC-I restricted antigenic epitope provides a more efficient immune protection than vectors expressing the full-length protein [21]. Therefore, we constructed an MCMV influenza vaccine vector by inserting the coding sequence for the H-2K^d^ MHC-I restricted peptide **IYSTVASSL** from the hemagglutinin (HA) of the H1N1 (PR8) IAV [34] into the C-terminus of the MCMV *ie2* gene. The resulting recombinant virus was called MCMV^IVL^ (Fig 1A). To test if the recombinant virus retained its capacity to replicate in host cells, virus replication was assessed by multi-step growth kinetics assays in NIH-3T3 cells *in vitro* and by *ex vivo* quantification of virus titers in livers, lungs and spleens 5 days post-infection (dpi) and in salivary glands 21 dpi. MCMV^IVL^ showed identical replication properties as the MCMV^WT^, both *in vitro* (Fig 1B) and *in vivo* (Fig 1C), indicating that the insertion of the IVL**_533-541_** epitope does not impair virus replication and spread.

**Fig 1.**
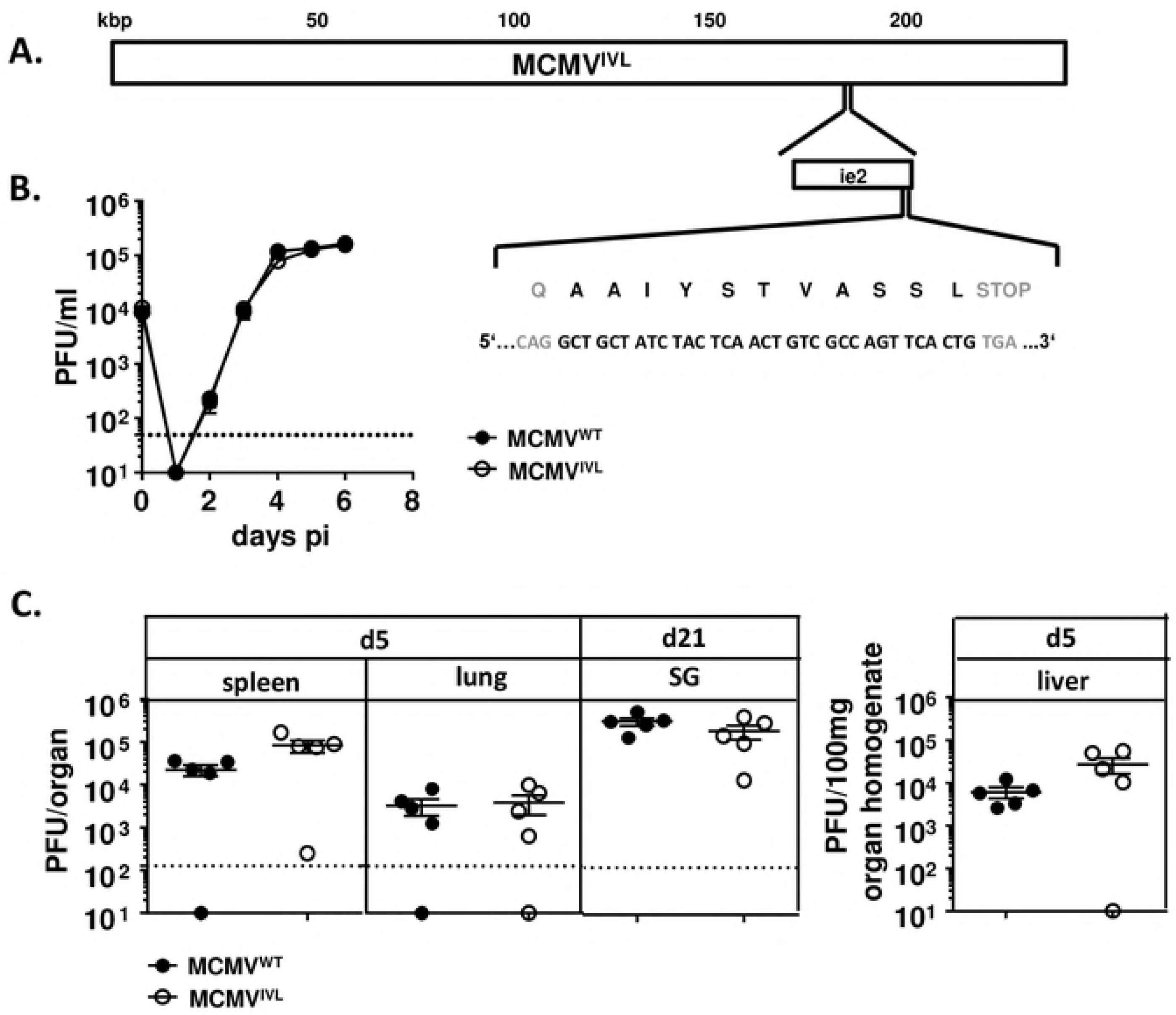
Generation of the recombinant MCMV expressing the _533_IYSTVASSL_541_ epitope. The sequence of IAV epitope **_533_**AAIYSTVASSL**_541_ (**IVL**_533-541_)** was introduced at the C-terminus of the MCMV *ie2* gene, and the growth of the recombinant virus MCMV^IVL^ was tested *in vitro* and *in vivo*. (A) The location of the *ie2* gene within the MCMV genome at ∼186-187kb position is shown; the insertion site of the peptide IVL**_533-541_** and the corresponding nucleotide sequences are magnified. (B) MCMV^IVL^ and wild-type MCMV (MCMV^WT^) growth at a multiplicity of infection of 0.1 was compared in NIH-3T3 cells. Virus titers in supernatants expressed as plaque-forming units (PFU) were established at indicated time points. Group means +/- standard error of the mean (SEM) are shown. The dashed line indicates the limit of detection. (C) BALB/c mice were i.p. infected with 2 x 10^5^ PFU of MCMV^IVL^ or MCMV^WT^ virus. Spleen, lung and liver homogenates were assayed for virus titers 5 days post-infection (dpi). Salivary-gland (SG) homogenates were assayed 21 dpi. Each symbol represents one mouse. Group means +/- standard error of the mean (SEM) are shown. The dashed line indicates the limit of detection.

### Intranasal immunization with MCMV^IVL^ induces a lower magnitude of CD8^+^ T cell response than intraperitoneal immunization

We have shown that mucosal infection with MCMV by the i.n. route induces memory inflation, although to a lower extent than upon the i.p. infection route [31]. To define if this pattern would hold true for the artificially incorporated influenza epitope as well, we compared the magnitude of the CD8^+^ T cell responses to MCMV^IVL^ and MCMV^WT^ induced via the i.n. and i.p. route, respectively. The kinetics of antigen-specific CD8^+^ T cell responses in peripheral blood was determined by IVL-tetramer staining. While we did not observe any difference at early times post immunization, i.p. immunization induced an overall stronger inflationary CD8^+^ T cell response during latency (Figs 2A and 2B). This pattern was observed both in relative terms (Fig 2A) and in absolute cell counts (Fig 2B). We next analyzed the IVL-specific CD8^+^ T cell responses in spleens, lungs and mediastinal lymph nodes (mLNs) at times of latency (>3months post infection (p.i)). Similarly, i.p. immunization induced higher levels of IVL-specific CD8^+^ T cells than the i.n. immunization in the spleen and lungs, both in relative (Figs 2C and 2D) and/or in absolute terms (Figs 2F and 2G). There were no significant differences in the mLNs (Figs 2E and 2H). We also analyzed the primed (CD11a^hi^CD44^+^) and IVL-specific CD8^+^ T cells during latency post-immunization for KLRG1 and CD62L expression in order to assess the fractions of central memory (T_CM_: KLRG1^-^CD62L^+^); effector (T_EFF_: KLRG1^+^CD62L^-^) and effector memory (T_EM_: KLRG1^-^CD62L^-^) subsets in blood, spleen and lung cells of i.p. and i.n immunized mice (S1 Fig). While the fraction of CD8T_CM_ cells remained relatively low in all compartments irrespectively of the route of administration, i.p. infection resulted in a response polarized towards effector cells, whereas i.n. immunization induced a higher fraction of EM cells in all analyzed organs. In sum, mucosal (i.n.) immunization with MCMV^IVL^ induces a systemic inflationary IVL-specific CD8^+^ T cell response, whereas the overall magnitude of the IAV-specific CD8^+^ T cell response is less pronounced compared to that induced in i.p. immunized mice. Moreover, CD8^+^ T cell responses induced via the mucosal route skew towards an effector memory phenotype.

### Intranasal immunization with MCMV^IVL^ facilitates the elimination of IAV in a CD8^+^ T cell-dependent manner

To test whether immunization with MCMV^IVL^ protects against IAV infection, latently immunized BALB/c mice were i.n. challenged with IAV. IAV titers in the lungs were quantified 5 dpi. Viral loads in mice that were immunized with MCMV^WT^ via either the i.p. or the i.n. route were comparable to those detected in mock-immunized mice (Fig 3A). In contrast, mice immunized with MCMV^IVL^ via the i.n. route showed significantly lower IAV titers than in any other group. Interestingly, i.p. MCMV^IVL^ immunization also resulted in reduced IAV loads, but to a lower extent than the i.n. immunization (Fig 3A). Similarly, animals immunized with MCMV^WT^ suffered the most severe weight loss whilst i.n. immunization of MCMV^IVL^ led to the least pronounced body weight loss. I.p. immunization with MCMV^IVL^ displayed an intermediate level (Fig 3B). Numerous studies have reported that CD8^+^ T cells play an important role in protecting against influenza infection [35, 36] and it was reasonable to assume that our vector provided immune protection by eliciting CD8^+^ T cell responses. To formally prove that efficient immune control of IAV observed in the MCMV^IVL^ (i.n.) immunized group depends on CD8^+^ T cells, we depleted these cells by systemic treatment of mice with a depleting anti-CD8α antibody (depletion efficiency is shown in S2A Fig) one day prior to IAV challenge and quantified viral titers in lungs 6 dpi. While the virus titer was below the detection limit in mice that were i.n. immunized with MCMV^IVL^ and received isotype control antibodies, CD8^+^ T cell depletion indeed resulted in a significant increase of the IAV titer to levels comparable to the groups that were i.p. immunized with MCMV^IVL^ and to control mice immunized with MCMV^WT^ by i.n. route (Fig 3C). CD8^+^ T cell depletion also slightly increased virus titers in both control groups - MCMV^IVL^ (i.p.) and MCMV^WT^ (i.n.), but not as pronounced as in the MCMV^IVL^ i.n. immunized group (Fig 3C). Similar as Fig. 3B, animals of all experimental groups showed a comparable body weight loss post-challenge, whereas i.n. MCMV^IVL^ immunized mice showed a faster recovery than the i.p. immunized mice (Fig 3D). Of note, this difference disappeared in the groups lacking CD8^+^ T cells (Fig 3E). Together, these data demonstrate that IAV-specific CD8^+^ T cells induced by the mucosal (i.n.) administration of MCMV^IVL^ confer protection against IAV in the lungs of vaccinated mice.

We further compared the lung pathology upon IAV challenge by histology. A moderate perivascular inflammation was observed in the lungs of most mice (stars), and to a lesser degree surrounding the bronchioles (arrows) (Fig 3F). The CD8^+^ T cell depleted group showed more severe pathology than isotype-treated controls, but the difference was not very pronounced (Fig 3G). Taken together, these data imply that intranasal immunization with the MCMV^IVL^ vector can limit IAV growth in the lungs by inducing IAV-specific CD8^+^ T cell responses, whereas the clinical outcome is only moderately improved.

### Intranasal immunization with MCMV^IVL^ induces antigen-specific tissue-resident memory CD8^+^ T (CD8T_RM_) cells in the lungs

We assumed that intranasal MCMV^IVL^ immunization may control the IAV replication by inducing CD8T_RM_ cell responses in the lungs. In order to test this hypothesis we identified the CD8T_RM_ cell compartment by staining cells with the CD69 [22] and the CD103 [26] marker at times of MCMV latency (> 3 months p.i.), as shown in Fig 4A. Only a few CD8T_RM_ (CD69^+^CD103^+^) cells could be detected in the spleen and blood regardless of the route of immunization (Fig 4B). CD8T_RM_ cells frequencies and counts were significantly higher in the lungs of mice that were i.n. immunized with MCMV^IVL^ compared to the i.p. route (Figs 4B and 4C). Resident memory T cells reside in the mucosal tissue layer and are non-migratory [27]. To confirm that CD8T_RM_ cells induced in the lungs of mice i.n. immunized with MCMV^IVL^ are indeed located within the airway mucosa, *in vivo* cell labeling experiments were performed [37, 38]. Here, intravenous (i.v.) injection of a fluorescent anti-CD45 antibody prior sampling allows for the discrimination of circulating leukocytes (fluorescence-positive) from emigrated or tissue-resident leukocytes (fluorescence-negative). The CD69^+^CD103^+^ cells from lungs were virtually absent from the CD45-labelled fraction (S3 Fig). IVL-specific CD8T_RM_ fractions were present solely in the lungs but not in the spleen and the blood (Fig 4D) and few IVL-tetramer^+^ CD8T_RM_ cells were induced by MCMV^IVL^ i.p. immunization (Figs 4D-4F). In contrast, in mice immunized with MCMV^IVL^ via the i.n. route, approximately half of the CD8T_RM_ cells were IVL-tetramer^+^ (Fig 4F). CD8T_RM_ cells were also induced in the group that was i.n. immunized with MCMV^WT^, but not IVL-specific (S4A Fig). In addition, there was an overall higher percentage of the CD69^+^CD103^-^CD8^+^ T cells in the lungs of i.n. immunized mice (S4B Fig), although the absolute cell numbers did not show a significant difference (S4C Fig). CD8T_RM_ cells showed low expression of Eomes whereas CD69^+^CD103^-^CD8^+^ T cells showed high expression of Eomes which is consistent with primed CD8^+^ T cells (S4D Fig). In summary, i.n. immunization with MCMV^IVL^ induces an accumulation of IAV-specific CD8T_RM_ response in the lungs.

### Pulmonary CD8T_RM_ cells improve viral clearance and the production of CD8^+^ T cell-recruiting chemokines during IAV infection

Resident memory T cells reside in the epithelial barrier of mucosal tissue [27] that is in close proximity to the airways. Hence, they can respond rapidly to a virus challenge at the site of infection [25, 27]. To define the relevance of lung CD8T_RM_ cells in protection against IAV challenge, we specifically depleted the airways CD8^+^ T cells by i.n. administration of αCD8 antibodies one day before challenge (Fig 5A, S2B and S2C Figs). To assess the impact of pulmonary CD8^+^ T-cell depletion on antiviral immunity, on day 4 post-challenge, we quantified the IAV titers in the lungs of i.n. MCMV^IVL^ immunized mice. Strikingly, targeted depletion of pulmonary CD8T_RM_ cells was associated with a significantly higher viral burden during IAV infection (Fig 5B). The data indicate that CD8T_RM_ cells induced by i.n. immunization with MCMV^IVL^ promote the clearance of IAV.

Influenza virus infection can induce a vigorous cytokine storm in airways and lungs, which promotes the recruitment of inflammatory cell. IFNγ as a pivotal antiviral cytokine is expressed early after influenza virus infection [39]. It has been demonstrated that CD8T_RM_ cells activate rapidly when they re-encounter the cognate antigen and provide protection by secreting cytokines such as IFNγ and granzyme B [40, 41]. *Morabito et al.* showed that intranasal immunization with an MCMV-based vaccine vector induced CD8T_RM_ cells and IFNγ was secreted at the very early time upon challenge during RSV infection [25].

Therefore, we measured the production of IFNγ in the bronchoalveolar lavage fluid (BALF) early upon challenge. IFNγ levels were generally low at day 2 post-challenge and no difference could be observed between groups regardless of airway CD8^+^ T cells depletion (Fig 5C). On day 4, the IFNγ level was significantly increased compared to the level on day 2, but more IFNγ was induced in the control group than in the one lacking airway CD8^+^ T cell in the lungs (Fig 5C). IFNγ was also induced in the MCMV^IVL^ i.p. immunization group and extremely low level of IFNγ could be detected in the MCMV^WT^ control group (S5A Fig). Thereby suggesting that primary cognate antigen immunization is needed for the rapid IFNγ secretion and that resident CD8^+^ T cells may not be the major IFNγ producer. In contrast to these effects, depletion of lung airway CD8^+^ T cells increased the concentration of IL-6 as compared to the group that was intranasally immunized with MCMV^IVL^ and treated with isotype control antibodies (Fig 5D). Similarly, a higher concentration of IL-6 was also detected in the i.p. immunization group, whereas the MCMV^WT^ control group displayed the highest IL-6 levels (S5B Fig). Finally, very low levels of other cytokines could be detected in all groups, both on day 2 and 4 post-challenge and regardless of the depletion of the airway CD8^+^ T cell (S5C Fig), suggesting that the presence of pulmonary CD8T_RM_ cells does not affect the Th1, Th2 and Th17 immune profile during early IAV infection.

It has been demonstrated that T_RM_ cells help to recruit immune cells to the infection site through the induction of chemokines such as CCL3 and CXCL9 in the female reproductive tract (FRT), and CCL4 in the lungs, either by direct chemokine expression or by their induction in nearby cells, such as epithelial cells [24, 25].

To determine whether i.n. immunization with MCMV^IVL^ induced inflammatory chemokines expression upon IAV challenge, a series of inflammatory chemokines were measured in the BALF on day 2 and day 4 upon IAV challenge (Figs 5E and 5F). As shown in Fig 5E, airway depletion of CD8^+^ T cells reduced CCL3, CCL4, CCL5 levels on day 2. On day 4, CCL3 and CCL4 levels were significantly higher in the MCMV^IVL^ i.n. group than in the MCMV^IVL^ i.p. and in the MCMV^WT^ i.n. immunization groups. Airway CD8^+^ T cell depletion reduced the level of CCL3 and CCL4 to values in the i.p. MCMV^IVL^ immunization group (Fig 5F). CXCL9 levels were comparable between the MCMV^IVL^ i.n. and i.p. immunization groups, but dramatically lower in the MCMV^WT^ immunization group (Fig 5F), which was consistent with the low IFNγ level in the BALF, as IFNγ is known as an inducer of CXCL9, which then acts as a T cell-attracting chemokine. Together, these data indicate that CD8T_RM_ cells induced by i.n. immunization with MCMV^IVL^ promote the induction of the pro-inflammatory chemokines CCL3, CCL4, CCL5 and CXCL9, along with a reduction of IL-6 in the lungs.

### CD8T_RM_ cells facilitate the expansion of CD8^+^ T cells in the lungs

Since i.n. immunization induced stronger chemokine responses in comparison to the i.p. immunization route, we decided to define whether CD8T_RM_ cells induced by MCMV^IVL^ promoted the accumulation of CD8^+^ T cells in the lungs. CD8^+^ T cell numbers on day 2 and 4 post IAV challenge were quantified in presence or absence of airway CD8^+^ T cells. In the MCMV^IVL^ i.n. immunization group, IVL-specific and total CD8^+^ T cell numbers increased from day 2 to day 4 post-challenge, but only in mice that were not depleted for airway CD8^+^ T cells (Figs 6A and 6B). IVL-specific CD8^+^ T cell counts in the lung tissue and BAL were slightly higher in the MCMV^IVL^ i.n. group than in the i.p. immunized group (S6A and S6B Figs). Interestingly, this differed from results prior to IAV challenge, where significantly larger amounts of IVL-specific CD8^+^ T cells were detected in the i.p. group (Fig 2G). IVL-specific and total CD8^+^ T cell counts increased significantly in the BALF of i.n. immunized mice by day 4 post IAV challenge, indicating that CD8^+^ T cells accumulate in the lungs and migrate to the bronchoalveolar space (Figs 6C and 6D). In mice where CD8^+^ T cells were depleted prior to challenge, very few IVL-tetramer^+^ CD8^+^ T cells (Fig 6C) and CD8^+^ T cells (Fig 6D) could be detected, both on day 2 and at day 4 post-challenge.

CD8^+^ T cells in the lung tissue were further analyzed by *in vivo* labeling of circulating cells. Anti-CD45 antibodies were injected i.v. 3-5 min prior to euthanasia and organ collection. The IVL-specific CD8^+^ T cell population failed to expand upon airway CD8^+^ T cell depletion, with significantly lower numbers in CD45^-^ subset on day 4. IAV-specific CD8^+^ T cell counts showed an increased trend both in the CD45^+^ and in the CD45^-^ subsets on day 4 post-challenge (Fig 6E). Airway CD8^+^ T cell depletion prevented also the expansion of total CD8^+^ T cells counts on day 4 (Fig 6F). Interestingly, only the CD45 unlabeled fraction of the total CD8 pool expanded on day 4 (Fig 6F), in contrast to IVL-specific CD8^+^ T cells, where CD45^+^ cells also expanded (Fig 6E). It is important to note that the number of total CD45^-^ CD8^+^ T cells increased in the i.n. immunization group, arguing that the recruitment by CD8T_RM_ cells was antigen-independent. Thus, non-cognate antigen-specific cells were also attracted from the circulation to the lungs and this accumulation of cells was abrogated by mucosal CD8^+^ T cell depletion (Fig 6F).

We surmised that the accumulation of CD8^+^ T cells in the lungs and in the BALF might be due to an expansion of CD8T_RM_ upon IAV challenge. Surprisingly, the number of CD8T_RM_ cells in the lungs did not expand from day 2 to day 4; if anything, their frequency decreased (Figs 7A and 7B). Likewise, IVL-Tetramer^+^ CD8T_RM_ cell counts were also slightly reduced from day 2 to day 4 post-challenge (Fig 7C), although IVL-Tetramer^+^ CD8 counts in the BALF increased at the same time (Fig 6A). It is important to note that the effect of the i.n. depletion was local, since the frequencies and counts of IVL-specific CD8^+^ T cells in the blood (S6C and S6D Figs) and spleen (S6E and S6F Figs) did not significantly differ upon i.n. αCD8 or upon isotype-control antibody administration. Therefore, our data indicated that CD8T_RM_ cells may confer protection by recruiting circulating CD8^+^ T cells upon IAV challenge.

## Discussion

Influenza-specific CD8^+^ T cells are known to contribute to virus elimination, because the clearance of influenza virus is delayed in T cell-deficient mice [5, 42]. However, previous evidence did not clarify whether vaccines solely inducing influenza-specific CD8^+^ T cell responses improve immune protection. To avoid confounding humoral immune responses and focus on the potential of optimally primed CD8^+^ T cells in protecting against influenza, we generated a new MCMV based vaccine vector. CMV vaccine vectors expressing exogenous antigenic peptides fused to a CMV gene induce CD8^+^ T cell responses of unparalleled strength [13, 15, 21, 25]. We show here that robust CD8^+^ T cell responses against a single MHC-I restricted epitope derived from the HA protein of IAV, promote the clearance of IAV from lungs, but only upon i.n. immunization. While some pathology was observed even in immunized mice, arguing that the protection was not complete, depletion assays confirmed that CD8^+^ T cells are crucial for the immune protection observed in our model. Remarkably, immunization by the i.p. route induced even stronger systemic CD8^+^ T cell responses, but very poor protection. This conundrum was resolved once we noticed that only i.n. immunization induces T_RM_ responses in the lung.

CD8T_RM_ cells act as sentinels in the host and form the first line of defense, providing rapid and effective protection to fight against pathogens invasion [23, 25, 27, 43]. Prior studies have revealed that direct delivery of vaccines to the target tissue is necessary for generation of T_RM_ cells [25, 44] and that sustained lung CD8T_RM_ responses in MCMV-infected mice are generated by immunoproteasome-independent antigenic stimulation [45], akin to the CD8 expansions in memory inflation [21], arguing that they are induced by similar and overlapping mechanisms. This was consistent with our observation that approximately half of CD8T_RM_ cells induced by MCMV^IVL^ i.n. immunization were IVL-specific, arguing that site-specific immunization with MCMV is necessary for the generation of memory-inflation-like antigen-specific CD8T_RM_ cells. Some prior studied have claimed that skin-resident CD8T_RM_ cells may confer protection in an antigen-unspecific manner [46], whereas others argued that only the antigen-specific CD8T_RM_ cells respond to cognate antigens [47]. MCMV^WT^ induced robust CD8T_RM_ responses in our model, but these were not specific for IAV, and did not provide any protection against IAV in our study. Site-specific anti-CD8α antibody administration depleted CD8 T_RM_, and increased IAV titers in immunized mice, indicating that CD8T_RM_ cells facilitated IAV elimination. Thus, the protection against IAV challenge required antigen-specific CD8T_RM_ cells in our model.

Early upon IAV challenge, the IVL-specific CD8^+^ T cells expanded strongly and rapidly in the lungs of i.n. immunized mice. The non-specific CD8^+^ T cell population also expanded dramatically, suggesting that the expansion was not antigen driven. However, total and IVL-specific CD8^+^ T cells expanded poorly in the lungs when airway CD8^+^ T cells were depleted. Finally, IAV challenge expanded IVL-specific CD8^+^ T cell counts in the blood and spleen of i.n. immunized mice to levels observed in the i.p. immunized controls, although the levels were significantly lower in the i.n. immunization group before challenge. Overall, these results indicated that i.n. immunization facilitated CD8^+^ T cell responses upon challenge, both locally in the lungs and systemically in the blood and spleen.

It has long been assumed that CD8T_RM_ cells have poor proliferative capacity upon challenge. Previous work has demonstrated that airway CD8^+^ T cells fail to expand *in vivo* upon intratracheal transfer [32] and that CD8T_RM_ cells induced by MCMV infection display a limited proliferative capacity in salivary glands [48]. However, this is in contrast to two recent studies demonstrating that CD8T_RM_ cells in the skin [47] and FRT [49] maintain the capacity of *in situ* proliferation upon cognate antigen stimulation. Such stimulation differentiates circulating effector memory CD8^+^ T cells into CD8T_RM_ cells without displacing the pre-existing CD8T_RM_ population [47]. In our study, CD8^+^ T cells accumulated in the lungs upon IAV challenge, but the CD8T_RM_ population did not expand and the number of antigen-specific CD8T_RM_ cells even displayed a reduction trend. Therefore, our data argued that either lung CD8T_RM_ in general or CD8T_RM_ induced by MCMV i.n. immunization in particular may behave differently from CD8T_RM_ in other organs. This distinction, however, goes beyond the scope of our current work and remains to be addressed in future studies.

We have shown in this study that concentrations of CCL3, CCL4 and CXCL9 in the BALF of the MCMV^IVL^ i.n. immunization group are significantly higher than in MCMV^IVL^ i.p. or MCMV^WT^ i.n. immunized mice. Intravital CD45 labeling showed that CD8^+^ T cells accumulating in the lungs are sequestered from the bloodstream, but not CD8T_RM_, arguing that circulating antigen-specific cells were attracted into the lungs under the presence of mucosa-resident CD8^+^ T cells. This is in line with the work of *Schenkel* et al. showing a rapid local induction of chemokines CXCL9 and CCL3/4 in the FRT upon re-infection, and recruitment of memory CD8^+^ T cells from the periphery [24]. Depletion of mucosal CD8^+^ T cells depressed chemokine levels in the BALF to levels seen in the i.p. immunization group. This, together with the high levels of IFNγ in the MCMV^IVL^ i.n. immunization group and extremely low IFNγ in MCMV^WT^ i.n. immunization points to a putative model where antigen-specific re-stimulation induces IFNγ, which drives chemokine responses that recruit CD8^+^ T cells from the bloodstream to the lungs.

In summary, our data demonstrate that CD8T_RM_ cells promote the induction of chemokines, which help to drive the recruitment of IVL-specific CD8^+^ T cells and facilitates the elimination of IAV. Furthermore, optimal induction of CD8T_RM_ cells in the lungs by the MCMV vector can be only achieved after i.n. vaccination. Therefore, immunization with an MCMV vector at the local site provided CD8^+^ T cell-based protection against IAV infection. Our results therefore demonstrate that CD8^+^ T cell induction, and CD8T_RM_ in particular, contribute to vaccination outcomes in influenza infection independently of humoral immune responses, and the selection of the adequate immunization route plays a critical role in terms for promoting superior protective efficacy.

## Materials and Methods

### Ethics statement

Mice were housed and handled in agreement with good animal practice as defined by EU directive EU 2010/63 and ETS 123 and the national animal welfare body ‘‘Die Gesellschaft für Versuchstierkunde /Society of Laboratory Animals (GV-SOLAS)’’. Animal experiments were performed in accordance with the German animal protection law and were approved by the responsible state office (Lower Saxony State Office of Consumer Protection and Food Safety) under permit number: 33.9-42502-04-14/1709.

### Mice

BALB/c mice were purchased from Janvier (Le Genest Saint Isle, France) and housed in the animal facility of the HZI Braunschweig under SPF conditions according to FELASA recommendations [50].

### Cells

Bone marrow stromal cell line M2-10B4 (CRL-1972) and NIH-3T3 fibroblasts (CRL-1658) were purchased from American Type Culture Collection (ATCC). The cells were maintained in DMEM supplemented with 10% FCS, 1% L-glutamine, and 1% penicillin/streptomycin. C57BL/6 murine embryonic fibroblasts (MEFs) were prepared in-house and maintained as described previously [51].

### Viruses

BAC-derived wild-type murine cytomegalovirus (MCMV^WT^ clone: pSM3fr 3.3) [52] was propagated on M2-10B4 lysates and purified on a sucrose cushion as described previously [53]. Virus titers were determined on MEFs by plaque assay as shown elsewhere [54].

Recombinant MCMV was generated by the ‘‘en passant mutagenesis’’, essentially as described previously [55, 56]. In brief, we generated a construct containing an antibiotic resistance cassette coupled with the insertion sequence and the restriction site Sce-I. This construct was flanked by sequences homologous to the target region of insertion within the MCMV BAC genome. Then, the fragment containing the insertion sequences was integrated into the MCMV BAC genome by homologous recombination. In a second step, Sce-I was induced to linearize the BAC followed by a second round of induced homologous recombination to re-circularize it and select for clones that discarded the antibiotic selection marker but retained the inserted sequence.

The PR8M variant of Influenza A/PR/8/34 was obtained from the strain collection at the Institute of Molecular Virology, Muenster, Germany. Virus stocks from chorioallantoic fluid of embryonated chicken eggs were generated as previously described [57].

## Tetramers and Antibodies

**_533_IYSTVASSL_541_** (**IVL_533-541_**)-tetramer was bought from MBL (cat. NO.TS-M520-1), anti-CD8α depletion antibody (clone: YTS 169.4). Rat IgG2b isotype antibody (clone: LTF-2) was purchased from Bio X Cell. Antibodies for flow cytometry included anti-CD3-APC-eFluor780 (clone: 17A2; eBioscience), anti-CD4-Pacific Blue (clone: GK1.5; BioLegend), anti–CD8α-PerCP/Cy5.5 (clone: 53-6.7; BioLegend), anti-CD11a-PE/Cy7 (clone: 2D7; BD Bioscience), anti–CD44-Alexa Fluor 700 (clone: IM7; BioLegend), anti-CD45-APC-eFluor780 (clone:30-F11;Biolegend), anti-CD62L-eVolve 605 (clone: MEL-14; eBioscience), anti-CD127-PE & PE/Cy7 (clone: A7R34; BioLegend), anti-KLRG1-FITC & BV510 (clone: 2F1/KLRG1; BioLegend), anti-CD103-APC (clone: 2E7; BioLegend), anti-CD69-FITC (clone: H1.2F3; BioLegend) and anti-IFNγ-APC (clone: XMG1.2; BioLegend), anti-Eomes-PE & PE/Cy7 (clone: Dan11mag; eBioscience).

### Virus *in vitro* infection

NIH-3T3 cell monolayers were infected with MCMV^WT^ and MCMV^IVL^ at an MOI of 0.1, incubated at 37°C for 1h, upon which the inoculum was removed, cells were washed with PBS, and supplied with fresh medium. Cells were incubated for 6 days; the supernatant was harvested every day and stored at −80°C until titration.

### Virus *in vivo* infection

6 to 8 weeks old BALB/c female mice were infected with 2 x 10^5^ PFU MCMV^WT^ and MCMV^IVL^ diluted in PBS. For i.p. infection, 100 µl virus dilution was injected. For i.n. infection, mice were first anesthetized with ketamine (10 mg/ml) and xylazine (1 mg/ml) in 0.9% NaCl (100 μl/10 g body weight), then administered with 20 µl of virus suspension onto nostrils [31]. For IAV challenge, BALB/c mice that were latently (> 3 months) immunized with MCMV were i.n. inoculated with 220 focus forming units (FFU) or with 1100 FFU of PR8M influenza virus as described previously [31].

### Infectious virus quantification (MCMV)

MCMV virus from organ homogenates was titrated on MEFs with centrifugal enhancement as described previously [17].

### Infectious virus quantification (IAV)

Mice were sacrificed by CO_2_ inhalation, whole lungs were excised and mechanically homogenized using a tissue homogenizer. Tissue homogenates were spun down and supernatants were stored at −70°C. Lung virus titers were determined by using the focus-forming assay (FFA), as described before [57] with minor modifications. Briefly, MDCK cells were cultured in MEM, supplemented with 10% FCS, 1% penicillin/streptomycin. Supernatants of lung tissue homogenates were serially diluted in DMEM, supplemented with 0.1% BSA and N-acetylated trypsin (NAT; 2.5 µg/ml) and added to the MDCK cell monolayers. After 1h, cells were overlaid with DMEM supplemented with 1% Avicel, 0.1% BSA and NAT (2.5 µg/ml). After 24h cells were fixed with 4% PFA and incubated with quenching solution (PBS, 0.5% Triton X-100, 20 mM Glycin). Cells were then treated with blocking buffer (PBS, 1% BSA, 0.5% Tween^®^20). Focus forming spots were identified using primary polyclonal goat anti-H1N1 IgG (Virostat), secondary polyclonal rabbit anti-goat IgG conjugated with horseradish peroxidase (KPL) and TrueBlue^TM^ peroxidase substrate (KPL). Viral titers were calculated as FFU per ml of lung tissue homogenate.

### Isolation of lymphocytes from blood and organs

Blood, spleen and mLNs were prepared as described previously [31]. Lungs were perfused by injecting 5-10 ml PBS into the right heart ventricle. The lungs were cut into small pieces, resuspended in 1 ml RPMI1640 (0.5% FCS), and digested with 1 ml of RPMI1640 with DNase I (Sigma-Aldrich Chemie) and Collagenase I (ROCKLAND^TM^) in a shaker at 37°C for 30 min. Digested tissue was passed through cell strainers and single cell suspensions were washed with RPMI1640, centrifuged at 500x g for 10 min. Subsequently, the cells were resuspended in 7 ml of 40% Easycoll solution (Biochrom), overlayed onto 6 ml of 70% Easycoll solution in a 15 ml Falcon and centrifuged at 25 min at 1000x g at room temperature. The interface layer was transferred to a 5 ml tube, washed, and resuspended in RPMI1640 (10% FCS).

### Peptide stimulation

T cells were stimulated with peptides (1 µg/ml) in 85 µl RPMI 1640 for 1h at 37°C, supplemented with brefeldin A (10 µg/ml in 15 µl RPMI 1640) and incubated for additional 5h at 37°C. Cells incubated without any peptide in the same condition were used as negative controls. Cytokine responses were detected by intracellular cytokine staining.

### Cell surface staining, intracellular cytokine staining and flow cytometry

Blood cells and lymphocytes from spleen, lung and mLNs were stained with IVL**_533-541_**-tetramer-PE and surface antibodies for 30 min, washed with FACS buffer and analyzed. For intracellular cytokine stainings, the cells were first stained with cell surface antibodies for 30 min, washed and fixed with 100 µl IC fixation buffer (eBioscience) for 5 min at 4°C. Following this, cells were permeabilized for 3 min with 100 µl permeabilization buffer (eBioscience) and incubated with anti-IFNγ antibody for 30 min. Afterwards, cells were washed with FACS buffer and acquired using an LSR-Fortessa flow cytometer (BD Bioscience).

### *In vivo* cell labeling

Mice were intravenously (i.v.) injected with 3 µg anti-CD45-APC/eFluor780 (clone: 30-F11; BioLegend). Mice were euthanatized 3-5 min after injection, and blood, spleen and lungs were collected. Following their isolation from the respective compartment, lymphocytes were stained and analyzed as described above.

### *In vivo* CD8^+^ T cell depletion

For systemic *in vivo* CD8^+^ T cell depletion, published protocols [58, 59] were adapted as follows. BALB/c mice were i.p. injected with 200 µg anti-CD8α (αCD8: clone: YTS 169.4) or isotype antibody (Rat IgG2b: clone: LTF-2; Bio X Cell) one day before IAV challenge. To deplete mucosal CD8^+^ T cells in the lungs, BALB/c mice were i.n. administered 10 µg αCD8 or IgG2b in 20 µl of PBS one day before IAV challenge [36].

### Collection of bronchoalveolar lavage fluid (BALF)

Mice were sacrificed by CO_2_ inhalation, chest cavity was opened and skin and muscle around the neck were gently removed to expose the trachea. A catheter was inserted and lungs were carefully flushed with 1 mL PBS via the trachea. The BALF was transferred into a 1.5 ml tube and stored on ice. The BALF was centrifuged at 500x g at 4°C for 10 min. Supernatant was aliquoted and stored at −80°C until further analysis.

### Cytokine and chemokine quantification

Mouse IFNγ enzyme-linked immunosorbent assay (ELISA) MAX™ kits (BioLegend) and the bead-based immunoassay LEGENDplex™ Mouse Inflammation Panel (13-plex, BioLegend) were used to quantify IFNγ and other cytokine levels in the BALF according to the manufacturer’s instructions. The bead-based immunoassay LEGENDplex^TM^ Mouse Pro-inflammation Chemokine Panel (13-plex, BioLegend) was used to quantify multiple chemokine levels in the BALF.

## Histopathology

Lungs were harvested from BALB/c mice that were latently infected with MCMV^WT^ and MCMV^IVL^ and challenged with IAV during latency. Lungs were fixed in 4% formalin, paraffin embedded, sliced and hematoxylin and eosin (H&E) stained according to standard laboratory procedures.

## Statistics

One-way ANOVA analysis was used to compare multiple groups at single time points. Two-way ANOVA analysis was used to compare different groups at multiple time points. Comparisons between two groups were performed using Mann-Whitney U test (two-tailed). Statistical analysis was performed using GraphPad Prism 7.

## Acknowledgements

We wish to thank Inge Hollatz-Rangosch and Ayse Barut from the Cicin-Sain lab and Tatjana Hirsch from the group of Prof. Dr. Dunja Bruder for excellent technical assistance. We thank Ramon Arens for tetramer reagents and Prof. Dr. Dirk Busch for providing us streptamer reagents. This work was supported by the ERC Stg 260934 and the Helmholtz EU Partnering grant “MCMVaccine” to LCS, and by a Chinese Scholar Council fellowship to XZ.

## Figure legends

**Fig 2. Intranasal immunization of MCMV^IVL^ induces weaker CD8^+^ T cell response than intraperitoneal immunization.** BALB/c mice were infected with 2 x 10^5^ PFU MCMV^IVL^ via the i.n. or i.p. route or with MCMV^WT^ via the i.n. route. IVL-specific CD8^+^ T cell responses were analyzed by IVL-tetramers staining and flow-cytometry. (A, B) Blood leukocytes were analyzed at indicated time points upon infection to define the (A) percentage and (B) cell counts of IVL-specific (Tet^+^) CD8^+^ T cells in peripheral blood. Two independent experiments were performed and results were pooled and shown as group means +/- SEM (n=8-10). (E-H) IVL-specific CD8^+^ T cells in spleen, lungs and mLNs were quantified by IVL-tetramer staining 120 dpi as relative cell percentages (C, D, E) or absolute cell counts (F, G, H) per organ. Two independent experiments were performed and pooled data are shown. Each symbol represents one mouse, n=7-10. Group means +/- SEM are shown. Significance was assessed by Two-way ANOVA test (panels A, B) or one-way ANOVA test (Panels C-H). **\***P <0.05, **\*\***P<0.01, **\*\*\***P<0.001.

**Fig 3. Intranasal immunization with MCMV^IVL^ facilitates the elimination of IAV.** BALB/c mice were immunized with 2 x 10^5^ PFU MCMV^IVL^ or MCMV^WT^ via the i.n. or i.p. route. Mock controls received 100 µl PBS by i.p. route. Once latency was established (> 3 months p.i), mice were challenged with IAV (PR8). (A) IAV titers in the lungs on day 5 post-challenge (i.n., 220 FFU) by focus-forming assay (FFA). Two independent experiments were performed and pooled data are shown, n=10 (B) Body weight loss upon IAV challenge (i.n., 1100 FFU), n=3-7. (C) CD8^+^ T cells were depleted systemically by i.p. injection of 200 µg anti-CD8α antibody (αCD8) or isotype control antibody (IgG2b) one day before PR8 challenge (i.n., 1100 FFU). Virus loads in the lung homogenates were quantified on day 6 post-challenge by FFA. Each symbol represents one mouse, n=5. Group means +/- SEM are shown. (D, E) Body weight loss upon IAV challenge (i.n., 1100 FFU) without (D) or with (E) systemic CD8^+^ T cell depletion. Two independent experiments were performed and pooled data are shown, n=10. Group means +/-SEM are shown. (F) H&E staining of the lung tissue on day 5 post-challenge (i.n., 1100 FFU) with or without systemic depletion of CD8^+^ T cell. (G) The score of inflammation in the lungs upon IAV challenge (i.n., 1100 FFU). Bars indicate means, error bars are SEM. Significance was assessed by One-way ANOVA test or Two-way ANOVA test. **\***P<0.05, **\*\***P<0.01, **\*\*\*\***P<0.0001, ns: no significant difference.

**Fig 4. Intranasal immunization with the MCMV^IVL^ induces antigen-specific tissue resident CD8^+^ T cells in the lungs.** BALB/c mice were i.n. (∘) or i.p. (•) infected with 2 × 10^5^ PFU of MCMV^IVL^ virus. Leukocytes were isolated from peripheral blood, spleen and lungs at latency time (> 3 months p.i), stained with antibodies against CD3, CD4, CD8α, CD11a, CD44, CD103, CD69, IVL-tetramer and measured by flow cytometry. (A) Representative gating strategy to identify CD8T_RM_ cells (CD69^+^CD103^+^). (B) Percentage of CD8T_RM_ cells in the peripheral blood, spleen and lungs. (C) Absolute CD8T_RM_ cell counts in the lungs of i.n. and i.p. immunized mice. (D) Percentage and (E) absolute counts of IVL-specific CD8T_RM_ cells in the peripheral blood, spleen and lungs of the i.n. and i.p. immunization groups. (F) Representative gating strategy and percentage of IVL-specific cells within the CD8T_RM_ cell subset in blood, spleen and lungs of i.n. and i.p. immunized mice. Two independent experiments were performed and pooled data are shown. Each symbol represents one mouse, n=8-10. Group means +/- SEM are shown. Significance was assessed by One-way ANOVA (B, D, E, F) test or Mann-Whitney U test (C). **\*\***P<0.01, **\*\*\*\***P<0.0001.

**Fig 5. CD8T_RM_ cells facilitate the elimination of IAV.** BALB/c mice were i.n. or i.p. immunized with 2 x 10^5^ PFU MCMV^IVL^ or i.n. with MCMV^WT^. During latency (> 3 months p.i), mice were treated with αCD8 or IgG2b antibodies and challenged with IAV (PR8) (i.n., 1100 FFU). Leukocytes were isolated from lungs on day 4 post-challenge for flow cytometric analysis. (A) Graphic representation of the mucosal CD8^+^ T cell depletion protocol. (B) IAV titers in the lungs on day 4 post-challenge of MCMV^IVL^ i.n. immunized mice. Two independent experiments were performed and pooled data are shown. Each symbol represents one mouse, n=10. Group means +/- SEM are shown. (C-F) The concentration of inflammatory cytokines and chemokines were measured in the BALF on day 2 and day 4 post-IAV challenge. (C) The concentrations of IFNγ and (D) IL-6 in the BALF of each MCMV^IVL^ i.n. immunized mice are shown as symbols. Group means +/- SEM are shown. (E) The concentration of CCL3, CCL4, CCL5 and CXCL9 in the BALF on day 2 post-challenge. (F) The concentration of CCL3, CCL4 and CXCL9 in the BALF on day 4 post-challenge. Experiments were performed twice with 5-7 mice per group and representative data are shown. Bars indicate means, error bars are SEM. Significance was assessed by Mann-Whitney U test, One-way ANOVA test, or Two-way ANOVA test. **\***P<0.05, **\*\***P<0.01, **\*\*\***P<0.001, **\*\*\*\***P<0.0001.

**Fig 6. CD8T_RM_ cells facilitate the accumulation of CD8^+^ T cells in the lungs.**

BALB/c mice were immunized with 2 x 10^5^ PFU MCMV^IVL^ via the i.n. route. During latency (> 3 months p.i), mice were i.n. treated with αCD8 (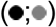) or IgG2b (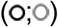) antibodies and challenged with IAV (PR8) (i.n., 1100 FFU) one day after. Leukocytes were isolated from lung tissue to analyze the CD8^+^ T cell response on day 2 and day 4 post-challenge. (A) Cell counts of IVL-specific CD8^+^ T cells in the lung tissue. (B) Cell counts of total CD8^+^ T cells in the lung tissue. (C) Cell counts of IVL-specific CD8^+^ T cells in the BAL. (D) Cell counts of total CD8^+^ T cells in the BAL. (E, F) anti-CD45 antibodies were injected intravenously 3-5 min before mice euthanasia and (E) IVL-specific CD8^+^ T cells or (F) Total CD8^+^ T cells that were intravitally labeled or remained unlabeled in the lung tissue were counted at 2 or 4 days post IAV challenge. Experiments were performed twice with 5-7 mice per group and representative data are shown. Each symbol represents one mouse, Group means +/- SEM are shown. Significance was assessed by One-way ANOVA test. **\***P <0.05, **\*\***P <0.01, **\*\*\***P <0.001, **\*\*\*\***P <0.0001.

**Fig 7. CD8T_RM_ cells do not expand upon IAV challenge.** BALB/c mice were immunized with 2 x 10^5^ PFU MCMV^IVL^ via the i.n. route. During latency (> 3 months p.i), mice were i.n. treated with αCD8 (•) or IgG2b (∘) antibodies and challenged with IAV (PR8) (i.n., 1100 FFU) one day after. Leukocytes were isolated from lung tissue to analyze the CD8^+^ T cell response on day 2 and day 4 post-challenge. (A) Percentage and (B) Counts of CD8T_RM_ cells in the lungs. (C) Counts of IVL-specific CD8T_RM_ cells in the lungs. Experiments were performed twice with 5-7 mice per group in total and representative data are shown. Each symbol represents one mouse. Group means +/- SEM are shown. Significance was assessed by One-way ANOVA test. **\***P <0.05, **\*\***P <0.01, ns: no significant difference.

## Supporting information

**S1 Fig. Phenotype of primed and IVL-specific CD8^+^ T cells.**

BALB/c mice were immunized with 2 x 10^5^ PFU MCMV^IVL^ via the i.p. or i.n. route. During latency (> 3 months p.i), leukocytes from blood, spleen and lungs were stained with cell surface markers CD3, CD4, CD8, CD11a, CD44, KLRG1, CD62L, IVL-tetramer and analyzed by flow cytometry. Primed cells are defined as CD11a^hi^CD44^+^), T_EFF_ as KLRG1^+^CD62L^-^, T_EM_ as KLRG1^-^CD62L^-^and T_CM_ as KLRG1^-^CD62L^+^. (A-B) The percentages of primed and IVL-tetramer^+^ CD8^+^ T cell subsets with different phenotypes in the blood (A & B), spleen (C & D) and lungs (E & F) are shown. Each symbol represents one mouse, pooled data from two independent experiments are shown (n=8-10). Group means +/- SEM are shown. Significance was assessed by Mann-Whitney U test (two-tailed). *P <0.05, **P <0.01, ***P <0.001, ****P <0.0001.

**S2 Fig. Efficiency of CD8^+^ T cell depletion.**

BALB/c mice were immunized with 2 × 10^5^ PFU MCMV^IVL^ by the i.n. route. (A) During latency (> 3 months p.i), mice were injected 200 µg αCD8 antibody (i.p.) to deplete total CD8^+^ T cells. Same amount of IgG2b antibody was given as isotype control. Leukocytes from blood, spleen and lung were analyzed by flow cytometry and representative flow cytometric panels in blood, spleen and lungs on day 1 post-depletion are shown. (B-C) Mice were administered with 10 µg αCD8 antibody (i.n.) to deplete airway CD8^+^ T cells in the lungs or IgG2b as a control. (B) The efficiency of mucosal CD8T_RM_ cell depletion in the lungs of MCMV^IVL^ i.n. immunized mice on day 1 post-depletion. (C) Total and IVL-specific CD8^+^ T cell counts in the blood of MCMV^IVL^ i.n. immunized mice on day 1 post airway CD8^+^ T cell depletion. Bars indicate means, error bars are SEM.

**S3 Fig. Efficiency of *in vivo* CD8^+^ T cell labeling.**

BALB/c mice were i.n. immunized with 2 x 10^5^ PFU MCMV^IVL^. During latency (> 3 months p.i), mice were i.v. injected 3 µg anti-CD45 antibody in 100 µl PBS 3-5min before euthanasia. Leukocytes were isolated from blood, spleen and lungs, stained with cell surface markers against CD4, CD8, CD69, CD103 before flow cytometry. (A) Representative dot plots of the cell surface expression of *ex vivo* labeled CD8 and *in vivo* labeled CD45. (B) Representative backgating showing that lung CD8T_RM_ (CD69^+^CD103^+^) cells are located exclusively within the CD45 unlabeled cells.

**S4 Fig. Mucosal immunization induces CD69^+^ and CD8T_RM_ cells, but MCMV^WT^ induces only IVL-unspecific CD8T_RM_.**

(A) BALB/c mice were immunized with 2 x 10^5^ PFU MCMV^WT^ via the i.n. route. During latency (> 3 months p.i), leukocytes were isolated from lungs, stained with cell surface markers against CD4, CD8, CD69, CD103 before flow cytometry. Representative dot plots of CD8T_RM_ and IVL-specific CD8T_RM_ cells. (B, C) BALB/c mice were immunized with 2 x 10^5^ PFU MCMV^IVL^ via the i.n. or i.p. route. The (B) percentage and (C) counts of CD69^+^CD103^-^CD8^+^ T cells immunized via the i.n. and i.p. route in the lungs. Pooled data from two independent experiments are shown. Each symbol represents one mouse, n=7-9. Group means +/- SEM are shown. (D) Eomes expression on different subsets of CD8^+^ T cells in the lungs. Significance was assessed by Mann-Whitney U test. **\*\*\***P <0.001, ns: no significance.

**S5 Fig. Inflammatory cytokines in the BALF upon IAV challenge.**

BALB/c mice were immunized with 2 x 10^5^ PFU MCMV^IVL^ via the i.n. or i.p. route or with MCMV^WT^ via the i.n. route. During latency (> 3 months p.i), MCMV^IVL^ (i.n.) immunized mice were administered with 10 µg αCD8 or 10 µg IgG2b antibody (i.n.). MCMV^IVL^ (i.p.) and MCMV^WT^ (i.n.) immunized mice were administered with 10 µg IgG2b antibody (i.n.). One day later, animals were challenged with IAV (PR8) (i.n., 1100 FFU). On day 2 and day 4 post-challenge, BALF was harvested and measured cytokines production by bio-plexing. The concentration of (A) IFNγ and (B) IL-6 in the BALF on day 4 post-challenge. Two independent experiments were performed and pooled data are shown. Each symbol represents one mouse, n=5-7. Group means +/- SEM are shown. (C) Cytokine concentrations in the BALF in different immunization group on day 2 and day 4 post-challenge. Bars indicate means, error bars are SEM. Experiments were performed twice with 5-7 mice each group. Significance was assessed by One-way ANOVA test. **\*\*\***P <0.001.

**S6 Fig. Mucosal immunization with MCMV^IVL^ induced vigorous CD8^+^ T cell responses in blood, spleen and lungs.**

BALB/c mice were immunized with 2 x 10^5^ PFU MCMV^IVL^ by the i.n. or i.p. route or with MCMV^WT^ by the i.n. route. During latency (> 3 months p.i), leukocytes were isolated from lungs. Mice were challenged with IAV (PR8) (i.n., 1100 FFU) one day after airway CD8^+^ T cell depletion. Leukocytes were isolated from lung, BAL, blood and spleen on day 4 post-challenge. (A) Count of IVL-specific CD8^+^ T cells in the lungs and (B) BAL. (C) Percentage of IVL-specific CD8^+^ T cells in the blood and (E) spleen. (D) Count of IVL-specific CD8^+^ T cell in the blood and (F) spleen. Two independent experiments were performed and pooled data are shown. Each symbol represents one mouse, n=5-7. Group means +/- SEM are shown. Significance was assessed by One-way ANOVA test. **\***P <0.05, **\*\***P <0.01.

## References

1. WHO. Influenza (Seasonal) Fact sheet N°211”. who.int.. 2014; Available from: http://www.who.int/en/news-room/fact-sheets/detail/influenza-(seasonal).

2. Barria, M.I., et al., Localized mucosal response to intranasal live attenuated influenza vaccine in adults. J Infect Dis, 2013. 207 (1): p. 115–24.

3. Ilyushina, N.A., et al., Live attenuated and inactivated influenza vaccines in children. J Infect Dis, 2015. 211 (3): p. 352–60.

4. Eichelberger, M., et al., Clearance of influenza virus respiratory infection in mice lacking class I major histocompatibility complex-restricted CD8+ T cells. J Exp Med, 1991. 174 (4): p. 875–80.

5. Bender, B.S., et al., Transgenic mice lacking class I major histocompatibility complex-restricted T cells have delayed viral clearance and increased mortality after influenza virus challenge. J Exp Med, 1992. 175 (4): p. 1143–5.

6. Yager, E.J., et al., Age-associated decline in T cell repertoire diversity leads to holes in the repertoire and impaired immunity to influenza virus. J Exp Med, 2008. 205 (3): p. 711–23.

7. Baumgarth, N. and A. Kelso, In vivo blockade of gamma interferon affects the influenza virus-induced humoral and the local cellular immune response in lung tissue. J Virol, 1996. 70 (7): p. 4411–8.

8. Topham, D.J., R.A. Tripp, and P.C. Doherty, CD8+ T cells clear influenza virus by perforin or Fas-dependent processes. J Immunol, 1997. 159 (11): p. 5197–200.

9. Sylwester, A.W., et al., Broadly targeted human cytomegalovirus-specific CD4+ and CD8+ T cells dominate the memory compartments of exposed subjects. J Exp Med, 2005. 202 (5): p. 673–85.

10. Holtappels, R., et al., Enrichment of immediate-early 1 (m123/pp89) peptide-specific CD8 T cells in a pulmonary CD62L(lo) memory-effector cell pool during latent murine cytomegalovirus infection of the lungs. J Virol, 2000. 74 (24): p. 11495–503.

11. Karrer, U., et al., Memory inflation: continuous accumulation of antiviral CD8+ T cells over time. J Immunol, 2003. 170 (4): p. 2022–9.

12. Munks, M.W., et al., Four distinct patterns of memory CD8 T cell responses to chronic murine cytomegalovirus infection. J Immunol, 2006. 177 (1): p. 450–8.

13. Karrer, U., et al., Expansion of protective CD8+ T-cell responses driven by recombinant cytomegaloviruses. J Virol, 2004. 78 (5): p. 2255–64.

14. Podlech, J., et al., Murine model of interstitial cytomegalovirus pneumonia in syngeneic bone marrow transplantation: persistence of protective pulmonary CD8-T-cell infiltrates after clearance of acute infection. J Virol, 2000. 74 (16): p. 7496–507.

15. Tsuda, Y., et al., A replicating cytomegalovirus-based vaccine encoding a single Ebola virus nucleoprotein CTL epitope confers protection against Ebola virus. PLoS Negl Trop Dis, 2011. 5 (8): p. e1275.

16. Hansen, S.G., et al., Profound early control of highly pathogenic SIV by an effector memory T-cell vaccine. Nature, 2011. 473 (7348): p. 523–7.

17. Dekhtiarenko, I., et al., The context of gene expression defines the immunodominance hierarchy of cytomegalovirus antigens. J Immunol, 2013. 190 (7): p. 3399–409.

18. Borkner, L., et al., Immune Protection by a Cytomegalovirus Vaccine Vector Expressing a Single Low-Avidity Epitope. J Immunol, 2017. 199 (5): p. 1737–1747.

19. Beverley, P.C., et al., A novel murine cytomegalovirus vaccine vector protects against Mycobacterium tuberculosis. J Immunol, 2014. 193 (5): p. 2306–16.

20. Klyushnenkova, E.N., et al., A cytomegalovirus-based vaccine expressing a single tumor-specific CD8+ T-cell epitope delays tumor growth in a murine model of prostate cancer. J Immunother, 2012. 35 (5): p. 390–9.

21. Dekhtiarenko, I., et al., Peptide Processing Is Critical for T-Cell Memory Inflation and May Be Optimized to Improve Immune Protection by CMV-Based Vaccine Vectors. PLoS Pathog, 2016. 12 (12): p. e1006072.

22. Gebhardt, T., et al., Memory T cells in nonlymphoid tissue that provide enhanced local immunity during infection with herpes simplex virus. Nat Immunol, 2009. 10 (5): p. 524–30.

23. Wakim, L.M., et al., The molecular signature of tissue resident memory CD8 T cells isolated from the brain. J Immunol, 2012. 189 (7): p. 3462–71.

24. Schenkel, J.M., et al., Sensing and alarm function of resident memory CD8(+) T cells. Nat Immunol, 2013. 14 (5): p. 509–13.

25. Morabito, K.M., et al., Intranasal administration of RSV antigen-expressing MCMV elicits robust tissue-resident effector and effector memory CD8+ T cells in the lung. Mucosal Immunol, 2017. 10 (2): p. 545–554.

26. Mackay, L.K., et al., The developmental pathway for CD103(+)CD8+ tissue-resident memory T cells of skin. Nat Immunol, 2013. 14 (12): p. 1294–301.

27. Jiang, X., et al., Skin infection generates non-migratory memory CD8+ T(RM) cells providing global skin immunity. Nature, 2012. 483 (7388): p. 227–31.

28. Lapuente, D., et al., IL-1beta as mucosal vaccine adjuvant: the specific induction of tissue-resident memory T cells improves the heterosubtypic immunity against influenza A viruses. Mucosal Immunol, 2018. 11 (4): p. 1265–1278.

29. Smith, C.J., et al., Murine CMV Infection Induces the Continuous Production of Mucosal Resident T Cells. Cell Rep, 2015. 13 (6): p. 1137–48.

30. Baumann, N.S., et al., Tissue maintenance of CMV-specific inflationary memory T cells by IL-15. PLoS Pathog, 2018. 14 (4): p. e1006993.

31. Oduro, J.D., et al., Murine cytomegalovirus (CMV) infection via the intranasal route offers a robust model of immunity upon mucosal CMV infection. J Gen Virol, 2016. 97 (1): p. 185–95.

32. McMaster, S.R., et al., Airway-Resident Memory CD8 T Cells Provide Antigen-Specific Protection against Respiratory Virus Challenge through Rapid IFN-gamma Production. J Immunol, 2015. 195 (1): p. 203–9.

33. Hombrink, P., et al., Programs for the persistence, vigilance and control of human CD8+ lung-resident memory T cells. Nat Immunol, 2016. 17 (12): p. 1467–1478.

34. Tamura, M., et al., Definition of amino acid residues on the epitope responsible for recognition by influenza A virus H1-specific, H2-specific, and H1- and H2-cross-reactive murine cytotoxic T-lymphocyte clones. J Virol, 1998. 72 (11): p. 9404–6.

35. Flynn, K.J., et al., Virus-specific CD8+ T cells in primary and secondary influenza pneumonia. Immunity, 1998. 8 (6): p. 683–91.

36. Slutter, B., et al., Lung airway-surveilling CXCR3(hi) memory CD8(+) T cells are critical for protection against influenza A virus. Immunity, 2013. 39 (5): p. 939–48.

37. Anderson, K.G., et al., Intravascular staining for discrimination of vascular and tissue leukocytes. Nat Protoc, 2014. 9 (1): p. 209–22.

38. Anderson, K.G., et al., Cutting edge: intravascular staining redefines lung CD8 T cell responses. J Immunol, 2012. 189 (6): p. 2702–6.

39. Samuel, C.E., Antiviral actions of interferon. Interferon-regulated cellular proteins and their surprisingly selective antiviral activities. Virology, 1991. 183 (1): p. 1–11.

40. Cheuk, S., et al., CD49a Expression Defines Tissue-Resident CD8(+) T Cells Poised for Cytotoxic Function in Human Skin. Immunity, 2017. 46 (2): p. 287–300.

41. Topham, D.J. and E.C. Reilly, Tissue-Resident Memory CD8(+) T Cells: From Phenotype to Function. Front Immunol, 2018. 9: p. 515.

42. Wells, M.A., P. Albrecht, and F.A. Ennis, Recovery from a viral respiratory infection. I. Influenza pneumonia in normal and T-deficient mice. J Immunol, 1981. 126 (3): p. 1036–41.

43. Pizzolla, A., et al., Resident memory CD8+ T cells in the upper respiratory tract prevent pulmonary influenza virus infection. Sci Immunol, 2017. 2 (12).

44. Zens, K.D., J.K. Chen, and D. L. Farber, Vaccine-generated lung tissue-resident memory T cells provide heterosubtypic protection to influenza infection. JCI Insight, 2016. 1 (10).

45. Morabito, K.M., et al., Memory Inflation Drives Tissue-Resident Memory CD8(+) T Cell Maintenance in the Lung After Intranasal Vaccination With Murine Cytomegalovirus. Front Immunol, 2018. 9: p. 1861.

46. Ariotti, S., et al., T cell memory. Skin-resident memory CD8(+) T cells trigger a state of tissue-wide pathogen alert. Science, 2014. 346 (6205): p. 101–5.

47. Park, S.L., et al., Local proliferation maintains a stable pool of tissue-resident memory T cells after antiviral recall responses. Nat Immunol, 2018. 19 (2): p. 183–191.

48. Thom, J.T., et al., The Salivary Gland Acts as a Sink for Tissue-Resident Memory CD8(+) T Cells, Facilitating Protection from Local Cytomegalovirus Infection. Cell Rep, 2015. 13 (6): p. 1125–1136.

49. Beura, L.K., et al., Intravital mucosal imaging of CD8(+) resident memory T cells shows tissue-autonomous recall responses that amplify secondary memory. Nat Immunol, 2018. 19 (2): p. 173–182.

50. rodents, F.w.g.o.r.o.g.f.h.m.o., et al., FELASA recommendations for the health monitoring of mouse, rat, hamster, guinea pig and rabbit colonies in breeding and experimental units. Lab Anim, 2014. 48 (3): p. 178–192.

51. Reddehase, M.J., J. Podlech, and N.K. Grzimek, Mouse models of cytomegalovirus latency: overview. J Clin Virol, 2002. 25 Suppl 2: p. S23–36.

52. Jordan, S., et al., Virus progeny of murine cytomegalovirus bacterial artificial chromosome pSM3fr show reduced growth in salivary Glands due to a fixed mutation of MCK-2. J Virol, 2011. 85 (19): p. 10346–53.

53. Dag, F., et al., Reversible silencing of cytomegalovirus genomes by type I interferon governs virus latency. PLoS Pathog, 2014. 10 (2): p. e1003962.

54. Cicin-Sain, L., J. Podlech, M. Messerle, M. J. Reddehase, and U. H. Koszinowski., Frequent coinfection of cells explains functional in vivo complementation between cytomegalovirus variants in the multiply infected host. Journal of Virology, 2005. 79: p. 9492–9502.

55. Tischer, B.K., G.A. Smith, and N. Osterrieder, En passant mutagenesis: a two step markerless red recombination system. Methods Mol Biol, 2010. 634: p. 421–30.

56. Dekhtiarenko, I., L. Cicin-Sain, and M. Messerle, Use of recombinant approaches to construct human cytomegalovirus mutants. Methods Mol Biol, 2014. 1119: p. 59–79.

57. Blazejewska, P., et al., Pathogenicity of different PR8 influenza A virus variants in mice is determined by both viral and host factors. Virology, 2011. 412 (1): p. 36–45.

58. Salem, M.L. and M.S. Hossain, In vivo acute depletion of CD8(+) T cells before murine cytomegalovirus infection upregulated innate antiviral activity of natural killer cells. Int J Immunopharmacol, 2000. 22 (9): p. 707–18.

59. Kruisbeek, A.M., In vivo depletion of CD4-and CD8-specific T cells. Curr Protoc Immunol, 2001. Chapter 4: p. Unit 4 1.

